# Relative infectivity of the SARS-CoV-2 Omicron variant in human alveolar cells

**DOI:** 10.1101/2022.04.13.486321

**Authors:** Taewoo Kim, Kyoung Il Min, Jeong-Sun Yang, Jun Won Kim, Junhyung Cho, Yun Ho Kim, Jeong Seok Lee, Young Tae Kim, Kyung-Chang Kim, Jeong Yeon Kim, Kwon Joong Na, Joo-Yeon Lee, Young Seok Ju

## Abstract

With the emergence of multiple highly transmissible SARS-CoV-2 variants during the recent pandemic, the comparison of their infectivity has become a substantially critical issue for public health. However, a direct assessment of these viral characteristics has been challenging due to the lack of appropriate experimental models and efficient methods. Here, we integrated human alveolar organoids and single-cell transcriptome sequencing techniques to facilitate the evaluation. In a proof-of-concept study using the assay with four highly transmissible SARS-CoV-2 variants, including GR (B.1.1.119), Alpha (B.1.1.7), Delta (B.1.617.2), and Omicron (BA.1), a rapid evaluation of the relative infectivity was possible. Our results demonstrate that the Omicron (BA.1) variant is 3-5-fold more infectious to human alveolar cells than the other SARS-CoV-2 variants at the early phase of infection. To our knowledge, this study provides the first direct measurement of the infectivity of the Omicron variant and new experimental procedures that can be applied for monitoring newly emerging viral variants.

## Introduction

During the global spread of the coronavirus disease 2019 (COVID-19), many novel SARS-CoV-2 variants of concern (VOC) have emerged, posing an increased risk to global public health and quarantine (Davies et al., 2021; Sheikh et al., 2021; Twohig et al., 2022). International communities, such as GISAID (Khare et al., 2021), PANGO (O’Toole et al., 2021), and Nextstrain (Hadfield et al., 2018), have been monitoring and assessing the evolution of SARS-CoV-2 using periodic genomic sequencing of viral samples. The sequencing results have identified a few major SARS-CoV-2 variants, including GR (B.1.1.119) with the D614G variant (Korber et al., 2020), Alpha (B.1.1.7) (first detected in UK), and Delta (B.1.617.2) (first detected in India) (WHO). In Nov 2021, the Omicron variant (BA.1), characterized with 32 mutations in the spike protein, emerged from South Africa and is currently the dominant variant in many countries (Fernandes et al., 2020).

To understand the functional impacts and pathological characteristics of each VOC, various approaches have been conducted including epidemiological studies (Hart et al., 2022), spike binding affinity assay (McCallum et al., 2021b, 2021a; Meng et al., 2022), experimental model studies (Hui et al., 2022; Meng et al., 2022; Suzuki et al., 2022), and genetically engineered virus comparison studies. The epidemiological studies illustrate characteristics of viral transmission and clinical severity, but their underlying cellular and molecular mechanisms cannot be investigated. The spike binding assay measures the affinity between the virus spike protein and human receptor, but its biological impact is cryptic. The experimental model studies, including animal models (Ulrich et al., 2022) and cell lines, represent the issue of tropism in viral infection (J. Kim et al., 2020). Often, genetically engineered viruses with a specific mutation of interest, rather than natural viral variants (e.g., D614G (GR, Alpha, Delta and Omicron) (Zhou et al., 2021), N501Y (Alpha and Omicron) (Liu et al., 2022) or P681R (Delta) (Liu et al., 2021; Saito et al., 2022)), are used in the infection study, but these engineered viruses may not reflect the full characteristics of natural VOCs.

Despite all these efforts, the direct measurement of the relative infectivity for multiple VOCs, particularly the impact of the natural virus on physiological human tissues, has not been investigated. To address this issue, we developed a rapid, fully-controlled assay system, in which (1) human alveolar type-2 cells in lung organoids are single-cell dissociated and (2) are exposed to a pool of SARS-CoV-2 variants of known concentration for infection (**Figure 1A**). Then, after an hours to days-long culture, the full-length transcriptomes of the infected cells are sequenced at the single-cell resolution (adopting the SMART-seq3 technique) (Hagemann-Jensen et al., 2020) to capture the viral genomic mutations. These mutations are used to identify which VOCs are responsible for an individual cell’s infection.

**Figure 1.**
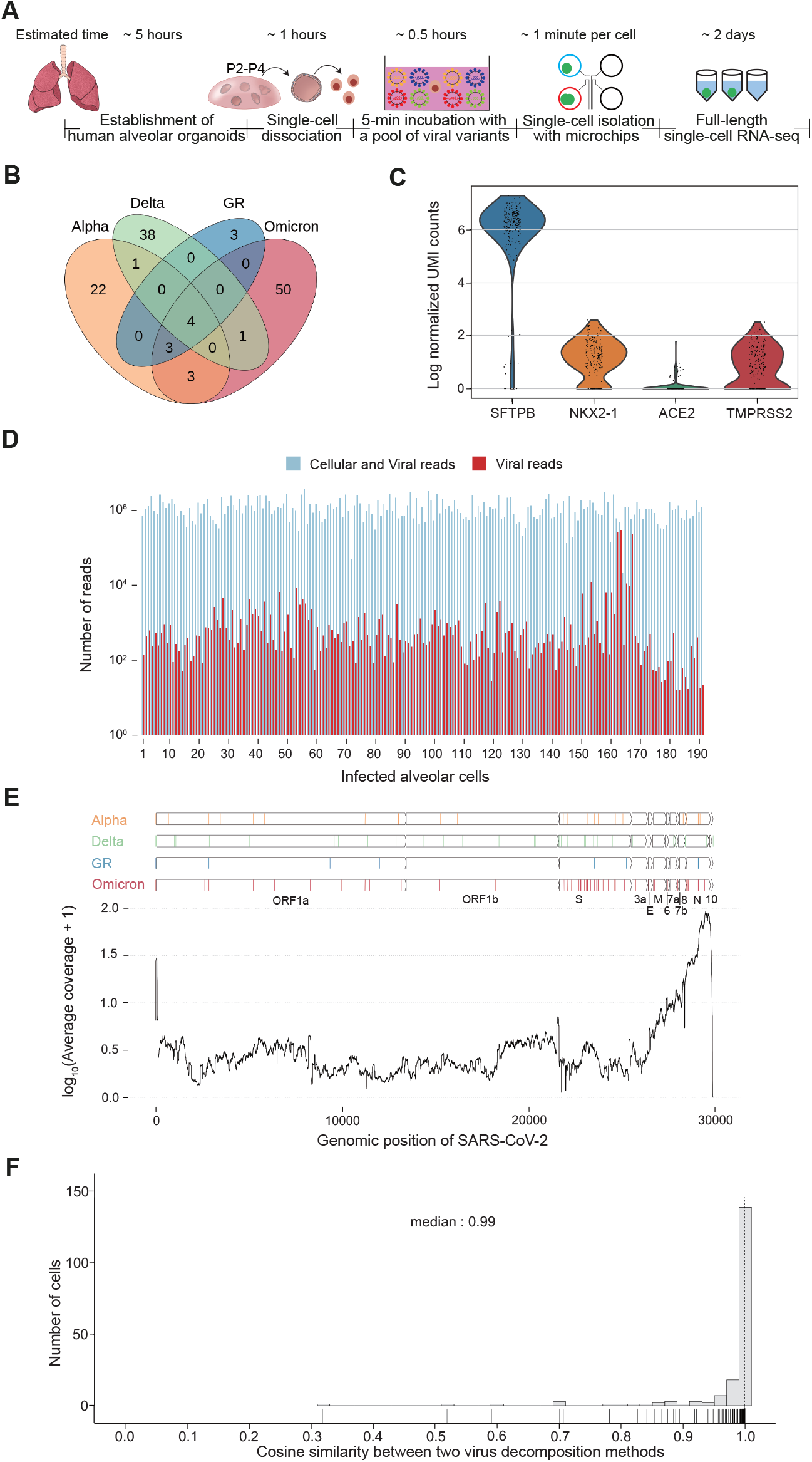
Integration of organoids and full-length single-cell RNA-seq techniques to understand the relative infectivity of SARS-CoV-2 variants. (A) Schematic diagram of the experimental procedures with expected experimental time. (B) Specific numbers of viral genomic mutations that are used for viral discrimination. (C) Expression levels of host genes (UMI counts) in the infected single-cells. (D) The number of reads from full-length single-cell transcriptome sequencing for each infected cell. Sky blue, total # of reads; Red, the number of viral reads. (E) Average coverage of viral transcripts. The genomic location of the viral mutations is shown on the top four panels. (F) Cosine similarity between two virus decomposition methods. Dashed line represents the median (0.99). Each cosine similarity was shown as a rug plot under histogram.

## Combined Results and Discussion

As a proof-of-concept study, we selected four SARS-CoV-2 variants, the GR clade virus (B.1.1.119), Alpha (B.1.1.7), Delta (B.1.617.2), and Omicron (BA.1), which are known as highly transmissible viruses during the pandemic. The viral particles were collected from Korean patients and maintained by the Korea Disease Control and Prevention Agency (KDCA). These viral variants can be differentiated by a combination of 125 genomic alterations (**Figure 1B).**

In our optimized infection experiments, viral incubation of alveolar cells was conducted at a multiplicity of infection (MOI) of 10 collectively, with each of the SARS-CoV-2 variants equally allocated for the incubation (**Methods**). On average, an alveolar cell interacted with 10 viral particles, and each variant had an equal chance of cellular infection. In our study, the viral incubation time was set mostly for five minutes, which turned out to be sufficient for the substantial infection of human alveolar type 2 (hAT2) cells. Then, the full-length transcriptome for SARS-CoV-2 infected single cells were sequenced by the SMART-seq3 technique with ~197Mb of sequencing throughput per cell, or ~1.3 M reads with 150 bp per cell, on average. The gene expression profiles of the host cells indicated that all of the infected single cells closely resembled alveolar type-2 cells (**Figure 1C**).

In the single-cell transcriptome of 191 infected cells that passed the quality check and threshold of the infection criteria (**Methods**), the viral sequences were observed in a substantial proportion ranging from 0.001% to 60% (**Figure 1D**). As previously reported (D. Kim et al., 2020), the 3’ genomic regions of the viral genome showed much higher RNA expression levels (**Figure 1E**). With the 125 viral genomic mutations as viral variant barcodes (**Figure 1B, Supplementary table 1**), we decomposed the fraction of each viral variant responsible for an individual cell’s infection using two independent algorithms (**Figure 1F; Methods**), which were overall concordant to each other. Of the 191 infected cells, 52, 85, and 52 cells were dominantly infected by single, double and multiple viral variants, respectively (27.2%, 44.5% and 27.2%), suggesting that multiple viral entry is possible in the experimental condition. For the remaining 2 cells (1.04%), unique viral variants could not be assigned due to the stochastic absence of the viral transcripts covering the mutation loci.

Despite the equal chance of infection, each SARS-CoV-2 variant showed strikingly different frequencies in the infected cells (**Figure 2A**). For instance, of the 52 cells with a single variant infection, 32 (61.5 %) were caused by the Omicron variant, followed by Alpha (n=18; 34.6 %), GR (n=1; 1.92%), and Delta (n=1; 1.92%). The Omicron variant was 2.46-fold more frequently observed than the random expectation (95% confidence interval = [1.88, 2.98]; p = 3.13×10^-9^), implying an ~4.8-fold higher infectivity than the other viruses by odds ratio under the same infectivity among viruses (**Methods**).

**Figure 2.**
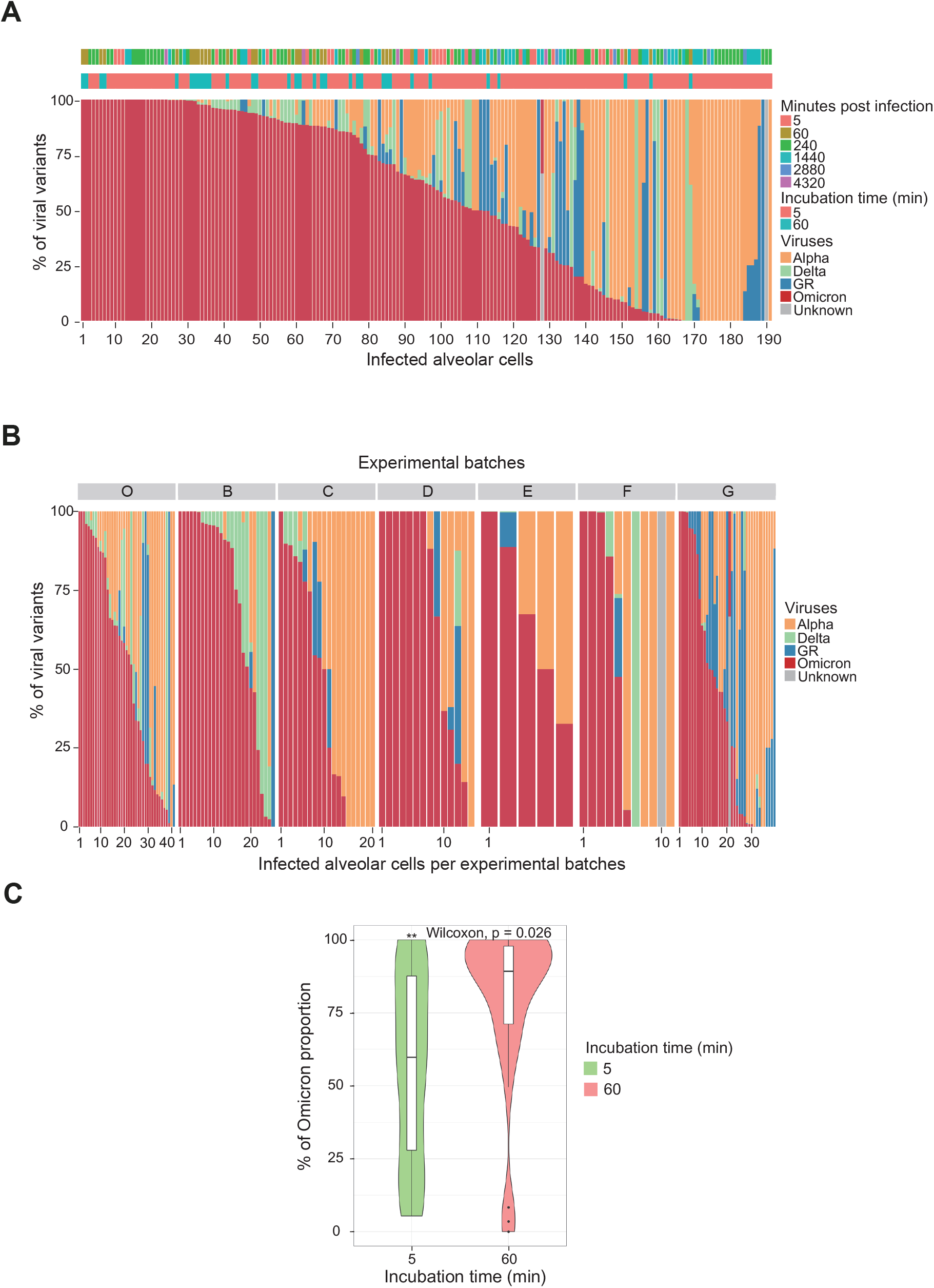
Higher infectivity of the Omicron variant. (A) The proportion of each viral variant in each infected cell. Experimental conditions (culture time post infection and viral incubation time) are shown on the top. The order of infected cells (X axis) is identical with the one in Fig 1D. (B) The proportion of each viral variant by 7 different experimental batches in this study. Only cells with a virus cell incubation time of 5 minutes are shown. (C) The proportion of the Omicron variant increases in the batch with a longer virus cell incubation time.

A similar conclusion was robustly drawn from a parallel analysis with all 191 cells, including the ones infected by two or more variants. Here, the Omicron variant was found in 161 cells (84.3%), followed by 106 (55.5%), 70 (36.6%) and 48 (25.1%) for the Alpha, Delta and GR variants, respectively, which is also biased toward the Omicron variant (p = 2.31×10^-35^) (**Figure 2A**). Taking into consideration the relative viral burden of each variant in an infected cell (a weighted average), the Omicron variant involved 105.9 cells (51.9%), outcompeting the other variants. This result means that the Omicron variant was observed 2.22-fold more frequently than the random expectation (95% confidence interval = [1.92, 2.50]; p = 5.76×10^-22^) and showed a 3.2-times higher infectivity than the other viruses. This result was concordant with the results drawn from the cells with the single variant infection. In the pairwise comparison with the other variants by odds ratio, the Omicron variant showed a 3.11 (against the Alpha), 18.7 (against the Delta) and 13.2 (against the GR) times higher infectivity in the assay. We believe that the dominance of the Omicron is robust because the trend was replicated in 7 independent batches (**Figure 2B**). Of note, in an experiment with a longer viral incubation time (60 min), the predominance of the Omicron variant was even higher (**Figure 2C**). However, it should be interpreted carefully due to the smaller number of infected cells in the specific experimental batch.

Recently, the higher infectivity of the Omicron than the other variants was shown in the upper airway (Hui et al., 2022; Meng et al., 2022; Suzuki et al., 2022). For alveolar cells, the Omicron was speculated to be less infective than the other variants, such as the Delta, because the Omicron depends more on an endocytic pathway for cellular entry rather than membrane fusion entry by TMPRSS2 (Meng et al., 2022). Our alveolar organoids express TMPRSS2 robustly as in the research by Meng et al. (**Figure 1B**), and our data directly indicate that the Omicron much more rapidly infects hAT2 cells than the other variants.

Of note, our data indicate a higher infectivity of the Alpha than the Delta (**Figure 2A**). An epidemiology study showed a higher hospital admission rate of the Delta (Twohig et al., 2022), but the spike binding affinity assay demonstrated enhanced affinities of the Alpha over Delta (Han et al., 2022). Presumably, the Alpha is more infective at the early phase of infection, but the Delta has a more serious impact in the late stage and/or in the clinical course. Our data should be interpreted carefully because four different viral variants competed in the series of experiments, and both the Alpha and Delta were all minor populations in the infected cells. To accurately measure the relative infectivity of the two variants specifically, another set of experiments, using the Alpha and Delta only, may be necessary.

One of the major advantages of our assay is that we can understand the infectivity of respiratory viruses against their target cell types, hAT2 cells. Our assay can be applied to any tissues whose organoids have a main target cell of a virus. Moreover, because the turn-around-time of our assay is ~3 days, the relative infectivity of viral variants can be quickly determined in a fully controlled condition. Compared with similar approaches based-on animal models (Ulrich et al., 2022) or 2D cell-lines (Peacock et al., 2022), our assay is free from the issue of viral tropism and recapitulates normal human tissue physiology better. The limitation of our method is the lack of immune cells, which have a role in viral progression *in vivo*.

Furthermore, our assay is able to decompose multiple viruses infecting a cell due to full-length single-cell RNA sequencing. To be specific, 10X Chromium cannot capture most of the viral genomic mutations. Technically, the manual picking of the infected single-cells is the rate limiting step in our assay. However, the process can be readily scalable by using a fluorescence-activated cell sorter (Hagemann-Jensen et al., 2021). Although this study mainly focused on the RNA transcripts from the virus, future studies may explore transcriptome changes of human genes at the single-cell resolution. To do so, infected cells should be incubated for a longer time, ideally for at least 24 h, as such a duration is necessary for alveolar cells to reprogram its transcription against viral infection (D. Kim et al., 2020; Youk et al., 2020).

To summarize, we demonstrated a new method that can be used to yield the relative infectivity of viral variants systematically and quickly. Our assay suggested that the Omicron variant is 3-5 times more infectious than the other SARS-CoV-2 variants against human alveolar cells. Together with other complementary approaches, our method will help to reveal the functional characteristics of emerging viral variants, especially for comparison among variants, in the future.

**Supplementary Table 1:**

A full list of genomic mutations of the Alpha, Delta, Omicron, and GR variants. (excel file)

## Materials and Methods

### Human tissues and alveolar organoid establishment

Human normal lung tissues were acquired from lung cancer patients with lobectomy surgery at SNUH with informed consent (IRB approval no. C-1809-137-975). From the human lung surgical samples, alveolar organoids were established as previously described (Youk et al., 2020). To remove cell-free RNA, we changed the RSPO-1 conditioned media to lyophilized RSPO-1 (80ng/ml) (R&D systems 4645-RS).

### Virus particle preparation for the competition assay

For the virus stock preparation, Vero cells were infected with a 0.01 MOI and grown in DMEM with 2% FBS and 1% P/S for 48 hours at 37 00B0;C with 5% CO2 as previously described (Youk et al., 2020). A purified viral particle stock was used to calculate the number of live viruses by Plaque-assay.

### Competition assay among VOCs: single cell infection and library preparation for single cells

First, human alveolar organoids were recovered by depolymerizing the Matrigel (Corning 354230) with Recovery solution (Corning 354253) at 4°C for 20 minutes. Furthermore, to remove the remaining Matrigel and dissociate the organoids into single cells, the organoids were incubated in Accutase (Stem Cell Technologies 07920) at 37°C for 5 minutes with additional mechanical pipetting. After washing, the cells were manually counted with the iNCYTO chip (iNCYTO DHC-N01). Cells were aliquoted into Protein LoBind 00AE; tubes (Eppendorf 0030108116) at 5,000 cells per tube. And then, each tube was incubated with different virus pool batches. The batches were recorded (**Figure 2B**). The cells were resuspended in 150 ul of Advanced DMEM/F12 (Thermo-fisher 12634010) with 1U/ml Penicillin/Streptomycin (Gibco 15630-080), 10mM HEPES (Gibco 15140-122), and 1% Glutamax (Gibco 35050-061) (v/v) (hereafter referred to as ADF+++).

Second, we prepared a total of 12,500 virus particles per virus. Every virus was mixed in a Protein LoBind^®^ tube to a final volume of 150 ul. After adding alveolar cells to the virus pool, the cells and viral particles were thoroughly mixed by pipetting for 20 times. The tubes were then incubated for 5 minutes at 37°C. Then, the cell-virus mix was washed for a total of 40,000X to completely remove the virus at the cell surface.

Fluorescence microscopy was used to increase the contrast of the live cells and decrease the single cell capture time. Furthermore, a microchip was used to isolate a single cell, and pictures were taken to store the cell status. The cells were incubated with 1000X CMFDA cell tracker (Thermo-Fisher C2925) for 10 minutes at 37°C. Next, the cells were loaded with 0.5% BSA (Sigma-Aldrich A8412) to decrease cell attachment on the smart aliquotor CE chip (iBioChip H2-SACE-5PK), and a single cell was picked up using 1 ul pipette after manual curation of the cell number using a fluorescence and phase contrast microscope.

For the SMART-seq3 library preparation, the minimal lysis buffer amount was first optimized for the cell in 1 ul PBS. It was determined that a 4 times volume before the 1st bead cleanup, compared to the original SMART-seq3 paper, was enough to successfully amplify the cDNA from the RNA. The lysis buffer was added to a captured cell, snap frozen, and stored at -80°C until lysis and reverse transcription. And then, the cells were thawed with the lysis buffer followed by the SMART-seq3 library protocol (Hagemann-Jensen et al., 2020). The pooled library was sequenced by Illumina NovaSeq by paired-end sequencing with an average of 190 Mb per cell.

### Viral genome mutations

To confirm the mutations of each virus that were used for the viral barcodes, we first conducted genome sequencing of the viral particles of each variant. Viral RNA was extracted by the QIAamp viral mini kit (Qiagen 52904) from a virus stock containing >10,000 viral particles. Then, a sequencing library was constructed with the NEBNext ARTIC SARS-CoV-2 FS kit (NEB E7658S). For each viral variant, paired-end sequencing was conducted by Illumina NovaSeq. Adapter sequences were removed with fastp (Chen et al., 2018. Then, the adapter-removed fastq files were aligned to the SARS-CoV-2 reference genome sequence, wuhCor1.fa (NC_045512v2) with BWA-MEM (Heng Li. arXiv:1303.3997). Primer sequences in the amplicons were removed using iVar (Grubaugh et al., 2019). Any human contamination reads were removed by Kraken (Wood et al., 2019). Then, the mutation sites were called using VarScan (Koboldt et al., 2012) and GATK HaplotypeCaller (Poplin et al., 2018). Candidate calls were manually inspected using Integrative Genomic Viewer (Thorvaldsdóttir et al., 2013), and in total, 125 clonal mutations (VAF >99%) were finally obtained (**Supplementary Table 1**) and compared among the four viral variants (**Figure 1B**).

### Data processing of the full-length single-cell transcriptome sequencings

From the pooled fastq file, we counted reads and UMIs of gene expression using zUMIs (Parekh1 et al., 2018). This condition is the same with the SMART-seq3 paper (Hagemann-Jensen et al., 2020). To align single-cell transcriptome sequences from infected cells, a joint reference genome sequence was established by concatenating the human genome (GRCh38.p13) and SARS-CoV-2 genome (wuhCor1.fa). A joint gene annotation file (gtf) was also generated by merging the primary annotation gtf of GRCh38.p13 and ncbiGenes.gtf downloaded from https://www.gencodegenes.org/human/ and https://hgdownload.soe.ucsc.edu/goldenPath/wuhCor1/bigZips/genes/, respectively. From the gtf file, we removed the nested exons of ORF1ab for calculating the gene expression of the viral transcripts. The UMI count matrix for exons was used for the transcriptome analysis.

For the quality control of the single cell RNA sequencing data, any cells with a mitochondria percentage above 40, number of expressed genes below 1,000, or total UMI count below 1,000 were excluded. The library size of each cell was adjusted by size factors calculated from the pooling-based deconvolution strategy using R package Scran (Lun et al., 2016). Library size adjusted cells were log normalized. Highly variable genes were selected by variance stabilizing transformation from R package Seurat (Stuart et al.,2019). Using the top 2,000 highly variable genes, a principal component analysis and UMAP embedding were calculated for visualization purposes. Single cell RNA sequencing analysis was done using Scanpy, if not otherwise stated (Wolf et al., 2018).

### Decomposition of the viral variants that infected a single-cell

We used the variant allele fraction (VAF) of each variant locus to calculate the proportion of the viral variant that infected a specific cell. Because not all 125 loci were covered by single-cell transcriptome sequencing, a specific consideration was necessary for an accurate decomposition. To this end, we used two different methods which were finally proven to be concordant to each other.

The first method is to utilize the average VAF of the multiple mutation loci. For each of the four viral variants, we calculated the average VAF of all covered unique mutation sites of a viral variant as a proxy of their fraction in a cell (F_alpha_, F_delta_, F_omicron_, and F_GR_). For cells with the sum of the average VAFs (F_sum_; F_alpha_+F_delta_+F_omicron_+F_GR_) smaller than 1, cellular infection was explained by an average of ~ 0.95 quite well. For most cells with F_sum_ > 1, we normalized each F value with the F_sum_ value. For cells explicitly infected but having F_alpha_+F_delta_+F_omicron_=0, F_GR_ was explicitly assigned to 1 because the GR variant has only 3 specific mutations (**Figure 1B**), and the viral variant is more likely to be unexposed in the transcriptome sequencing by chance. Lastly, for cells with more than two viral variants which are not covered for every unique mutation of that (NA), 1 - (F_sum_ without NA) is assigned as unknown

The second method is using nonnegative matrix factorization (NMF). Given a sample VAF matrix of size 125 × number of samples, the linear combination of the VOC genotypes that best explains each VAF column of the sample matrix was calculated. This process can be summarized as follows where the VOC matrix has a size of 125 × 4, and the weight matrix has a size of 4 × number of samples:

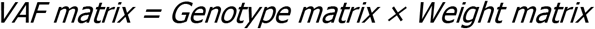

Because the two methods showed a high linear correlation, we used and showed the results obtained from the first method in **Figure 1F**.

### Criteria for infection at a single cells

In practice, some extracellular virus RNA may contribute to the viral reads in single-cell transcriptome sequencing although a cell is not infected. From empty wells in the single-cell transcriptome sequencing, we set a threshold for the viral infection as 3 or more viral UMI in more than 2 different viral genes.

### Virus transcripts average depth

The read-depth of deduplicated bam files was analyzed by SAMtools (Li et al., 2009) for each cell and averaged. We drew the gene plot above the coverage plot by gggenes (https://github.com/wilkox/gggenes).

### Statistical Analysis

To calculate the Omicron’s dominancy of the observed value against the expected value, for single and total infected cells, we adopted the proportion test with a random expectation of 0.25. Furthermore, to compare each variant’s infectivity, we calculated the odds ratios for each pair of viruses.

## Acknowledgements

We thank Myung Jin Yang (GSMSE KAIST) for the valuable technical help and comments. This project was supported in part by GENOME INSIGHT Inc. and the National Research Foundation of Korea (NRF-2020R1A3B2078973). This research was also supported by the Research Program 2020 funded by the Seoul National University College of Medicine Research Foundation (800-20200516)

